# Temperature and nutrient gradients correspond with lineage-specific microdiversification in the ubiquitous and abundant *Limnohabitans* freshwater genus

**DOI:** 10.1101/839407

**Authors:** Ruben Props, Vincent J. Denef

## Abstract

Most freshwater bacterial communities are characterized by a few dominant taxa, which are often ubiquitous across freshwater biomes worldwide. Our understanding of the genomic basis underlying this pattern is limited to a subset of taxa. Here, we investigated the genomic basis that enables *Limnohabitans*, a freshwater genus key in funneling carbon from primary producers to higher trophic levels, to achieve abundance and ubiquity. We reconstructed eight metagenome assembled genomes (MAGs) from this genus along broad environmental gradients existing in Lake Michigan, part of Earth’s largest surface freshwater system. *De novo* strain inference analysis resolved a total of 23 strains from these MAGs, which strongly partitioned into two habitat-specific clusters with co-occurring strains from different lineages. The largest number of strains belonged to the abundant LimB lineage, for which robust *in situ* strain-delineation had not previously been achieved. Our data show that temperature and nutrient levels may be two of the primary drivers of microdiversification within the *Limnohabitans* genus. Additionally, strains predominant in low and high phosphorus conditions had larger genomic divergence than strains abundant under different temperatures. Comparative genomics and gene expression analysis yielded evidence for the ability of LimB populations to exhibit cellular motility and chemotaxis, a phenotype not yet associated with cultured *Limnohabitans* strains. Our findings broaden historical marker gene-based surveys of *Limnohabitans* microdiversification, and provide *in situ* evidence of genome diversity and its functional implications across freshwater gradients.

## Importance

The importance of our work lies in the combined use of *in situ* genomic and gene expression data to deepen our understanding of what sustains the dominance of *Limnohabitans* bacterial populations in Lake Michigan, part of Earth’s largest surface freshwater system. We resolved for the first time the fine-scale genomic diversity within the LimB lineage, a lineage previously highlighted as invariant to environmental gradients. Our data shows that temperature variation can explain the observed geospatial distribution of the populations, and how adaptation to diverse nutrient regimes requires higher levels of genomic divergence. The gene expression profiles of this lineage further revealed the role of chemotaxis and motility in adapting to environmental conditions.

## Background

In natural and managed environments bacterial taxa partition across habitats at both coarse (1, 2) and fine taxonomic scales (e.g., > 97 – 99 % 16S rRNA gene similarity, or > ~96.5% genome-wide nucleotide identity) (3, 4). This genetic diversification, also labeled microdiversity, emerges from both mutational and gene gain/loss events (4–8) and represents an important trait-modifying process by which a bacterial taxon can achieve ubiquity, and thus maximize its niche coverage in an ecosystem (9). Usually, this microdiversity is masked by the consensus similarity thresholds used to define Operational Taxonomic Units (OTUs) in marker gene surveys, or hidden within consensus metagenome-assembled genomes (MAGs). In recent years several computational methods have been described to robustly infer microdiversity from 16S rRNA amplicon gene surveys (10, 11), as well as from metagenomic data (12–14). This has led to the numerous observations that small, or even no differences in 16S rRNA gene identity can lead to substantial alterations in, for example, optimal growth temperature (6), carbon substrate utilization (15), pH tolerance (16), and light preference (17). These trait differences are often reflected in the habitat partitioning across environmental gradients within the contiguous environment from which these taxa were sampled. Understanding how microdiversification enables bacterial taxa to adapt to changing environmental conditions can therefore facilitate our understanding and possibly mitigation of the short- and long-term impacts of global change (18, 19).

Freshwater lakes are known hotspots of global biogeochemical cycles and act as important environmental “sentinels” for local and global environmental change (20). This is also the case for Lake Michigan, which is part of the Laurentian Great Lakes system that contains an estimated 21% of the world’s surface freshwater. This ecosystem has been rapidly changing as a consequence of various anthropogenic stressors (21). The invasion of dreissenid mussels is causing drastic changes in the food web structure by decimating offshore primary production and indirectly causing nearshore harmful algal blooms (22–26). In addition to natural temperature and light gradients across seasons and depths, invasive mussel disturbances have led to steep estuary-to-offshore nutrient gradients across relatively small spatial scales.

Previous surveys of Lake Michigan have shown that the *Limnohabitans* genus, similar to many lake systems worldwide, and in particular the so-called “Lhab-A1 tribe” (27), is abundant, and that its abundance is largely unaffected by environmental gradients of temperature, trophic state, and light availability (28). *Limnohabitans* is a metabolically versatile, fast growing, morphologically diverse bacterioplankton genus that has been observed in nearly every lake system worldwide in high abundance (~12 %; (29)), and plays an important role in funneling carbon from primary producers to higher trophic levels (27, 30–34). The extensive isolate library of this genus has revealed a comprehensive metabolic versatility, ranging from photoheterotrophy (35) to putative ammonium and sulfur oxidation (36). In light of their biogeochemical importance, *Limnohabitans* is an excellent case study to understand how a taxon achieves abundance and ubiquity across environmental gradients. This knowledge can in turn be leveraged to better predict how their abundance and their functional contributions will respond to ongoing environmental changes.

Here, we studied microdiversification among core genes, differences in accessory genes, and *in situ* differential gene expression of populations belonging to the *Limnohabitans* genus across the environmental gradients of a well-studied Lake Michigan transect. For more than two decades this transect has been used to characterize the spatiotemporal changes in the pelagic food web and relate these changes to anthropogenic pressures and impacts of global change (37). It consists of three stations that sample the meso- to eutrophic Muskegon Lake freshwater estuary, the oligotrophic to mesotrophic nearshore (M15), and the ultra-oligotrophic offshore waters (M110). We hypothesized that depending on the magnitude of environmental changes microdiversity would be the predominant adaptation that governs the ability of *Limnohabitans* to maintain its prevalence and key role in the freshwater food web. To address this hypothesis we sampled the transect of Lake Michigan over large environmental gradients: temperature = 3 – 28 °C, depth = 1 – 110 m, PAR = 0 - 1042 W m^2^, total phosphorus = 3 – 30 μg L^−1^, chlorophyll *a* = 0.2 – 13 μg L^−1^. We used gene- and genome-centric metagenomics and metatranscriptomics to infer phylogeny, abundances, genome properties, gene expression, and inferred phenotypic traits (i.e. growth rate) of all putative *Limnohabitans* populations. We then performed variant detection and *de novo* strain inference on the reconstructed MAGs to evaluate the level of microdiversity within them.

## Results and Discussion

### Marker gene diversity

Full-length putative *Limnohabitans* 16S rRNA gene sequences were reconstructed from metagenomic data and incorporated in a phylogenetic tree together with publicly available isolate and environmental 16S rRNA gene sequences of the *Limnohabitans* genus (38). We reconstructed 45 unique sequences putatively classified as Limnohabitans of which 35 were phylogenetically placed within the *Limnohabitans* genus (**Figure 1**). The majority of sequences were assigned to the LimB lineage (n = 18) (38) and the Lhab-A4 tribe (n = 14) (27). This was in agreement with an earlier survey that found abundant LimB-related clone sequences in Lake Michigan (39), but is in striking contrast to the majority of isolate and clone reference sequences from other systems that belong primarily to the LimA, LimD and LimC lineages (38). This suggests that the LimB lineage may be difficult to cultivate using standard dilution-to-extinction cultivation methods. All reconstructed *Limnohabitans* sequences could be clustered into three OTUs (clustered at > 97 % similarity) which corresponds to the observations in a previous 16S rRNA gene amplicon survey (28). We found strong indications of microdiversity in the LimB and Lhab-A4 reconstructed sequences as within their respective clades the reconstructed sequences were on average 98.8 % similar, which is higher than the currently-adopted species threshold of 98.5% (40). Such a large number of highly-similar marker gene sequences have been observed before in several other aquatic genera (e.g., *Vibrio (41)*), of which not all the observed genotypes have been shown to clearly correspond to distinct environmental adaptations. Therefore, care must be taken to equate the number of reconstructed 16S rRNA gene sequences with the number of ecologically-cohesive populations in the environment. Interestingly, the sole sequence phylogenetically placed in the LimC lineage was 98.0 ± 0.6 % similar to those in the LimB lineage and was clustered together with the LimB sequences into a single OTU. These close similarities between lineages in the 16S rRNA gene have led other research groups to evaluate additional marker genes for delineating *Limnohabitans* taxonomic groups (32). Using this multi-marker approach, the majority of reported microdiversity has primarily been found in the LimC and LimA lineages, although the LimB lineage has been found to be less diverse but consistently the most abundant and ubiquitous (38, 42, 43). Jezberová et al. (2017) argue that the *Limnohabitans* lineage-specific probes may not be sensitive enough to delineate the microdiversity in the LimB lineage (43), suggesting that a genome-resolved approach may be necessary.

**Figure 1:**
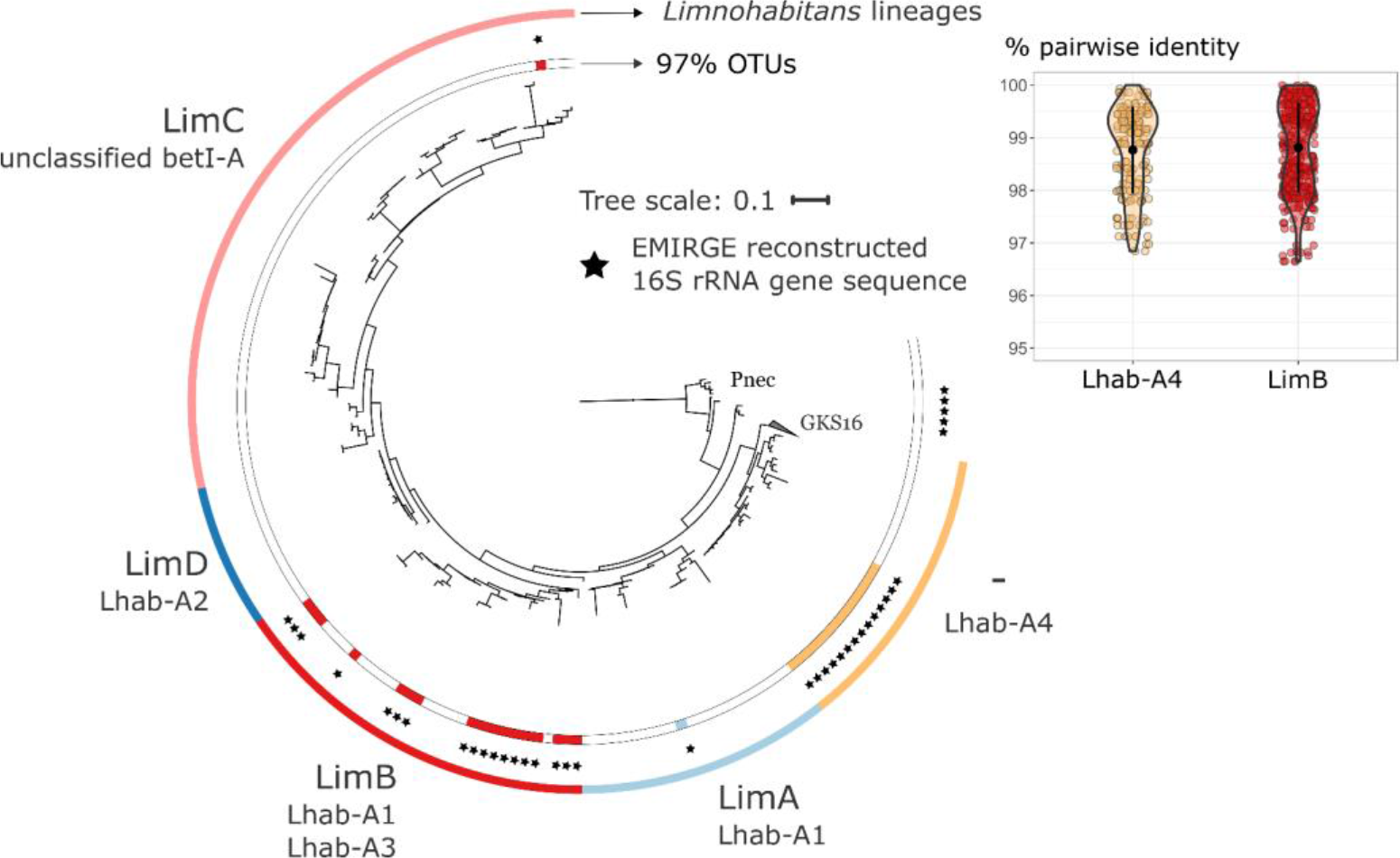
Phylogenetic tree of EMIRGE reconstructed 16S rRNA gene sequences and reference betaproteobacterium 16S rRNA gene sequences based on (38). Outer circle color coding indicate Limnohabitans lineages. Sequences within these lineages were also classified according to the Newton et al (2011) taxonomic framework into “Lhab” tribes. Inner color coding corresponds to the clustering of the EMIRGE reconstructed sequences at 97% average similarity. Pairwise % identity distributions are provided for the LimB and Lhab-A4 metagenome reconstructed 16S rRNA gene sequences. The tree was rooted in sequences of Enterobacter cancerogenus LMG 2693 (Z96078.1) and Escherichia vulneris (AF530476.1).

### Genomic diversity

#### Lake Michigan and Muskegon Lake contains diverse *Limnohabitans* taxa

We performed a genome-resolved study of the genetic diversity within the *Limnohabitans* taxa of Lake Michigan and its eutrophic estuary Muskegon Lake. Using a genome-centric metagenomic approach we were able to reconstruct 10 metagenome assembled genomes (MAGs) of medium-to-high estimated completeness (59 % – 97 %) and low estimated redundancy (< 3 %). Eight of these were confidently classified into the *Limnohabitans* genus using the MiGA taxonomic annotation pipeline (**Table 1**) (44). Several attempts to increase the quality of the medium-quality MAGs by modifying the assembly input (i.e. coverage-based normalization, randomized downsampling), assembly set-up (i.e. assembler software megahit/IDBA-UD, kmer range) or binning parameterization (i.e. minimum contig length, binning parameters) were not successful. The MAG coverage varied over more than an order of magnitude with MAG8.SU-M110-DCMD, further referred to as MAG8, being the overall best covered MAG. Most MAGs had a mean coverage < 10×, highlighting the need for deep sequencing to capture the rare *Limnohabitans* taxa. On average, more than 50% of each putative *Limnohabitans* population was replicating at the time of sampling as assessed by the index of replication (iRep > 1.5), an inferred measure for the replication state of the genome (45) (**see Fig. S1 posted at** https://doi.org/10.6084/m9.figshare.10265165). This suggests that both the abundant and rare populations were actively growing and thus metabolically active across the entire estuary to pelagic gradient.

**Table 1:**
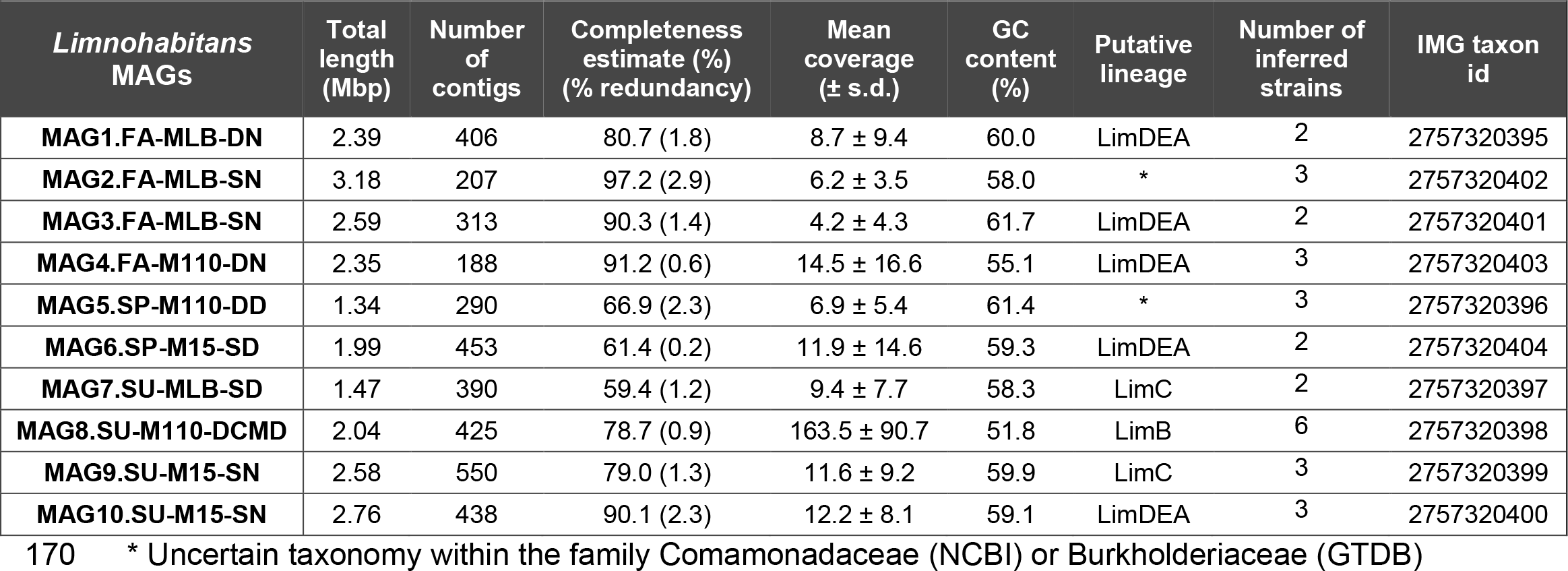
Assembly statistics and genome properties of the putative Limnohabitans MAGs described in this study. MAGs were labeled according to the environmental conditions of the sample from which they were reconstructed, putative lineages were based on placement in the phylogenomic tree (see Figure 2).

A phylogenomic tree based on 37 single-copy conserved core genes was made with the putative *Limnohabitans* MAGs and publicly available *Limnohabitans* isolate genomes (**Figure 2**). Two MAGs were placed into the LimC lineage and one in the LimB lineage (≤ 80 % ANI with reference genomes). The other seven were undefined as there were no reference genomes available for the LimDEA lineages, and two (i.e. MAG5 and MAG2) were part of a separate non-*Limnohabitans* taxonomic group that was not included in the tree. The LimB-placed MAG8 was the most abundant taxon while all other MAGs were comparable in mean abundance across the sampling stations (**Figure 2A**). LimB-associated marker gene sequences have been detected in many river and lake systems, and have been typically classified as “generalist” taxa as their (relative) abundance was reported to be invariant to environmental gradients (43).

**Figure 2:**
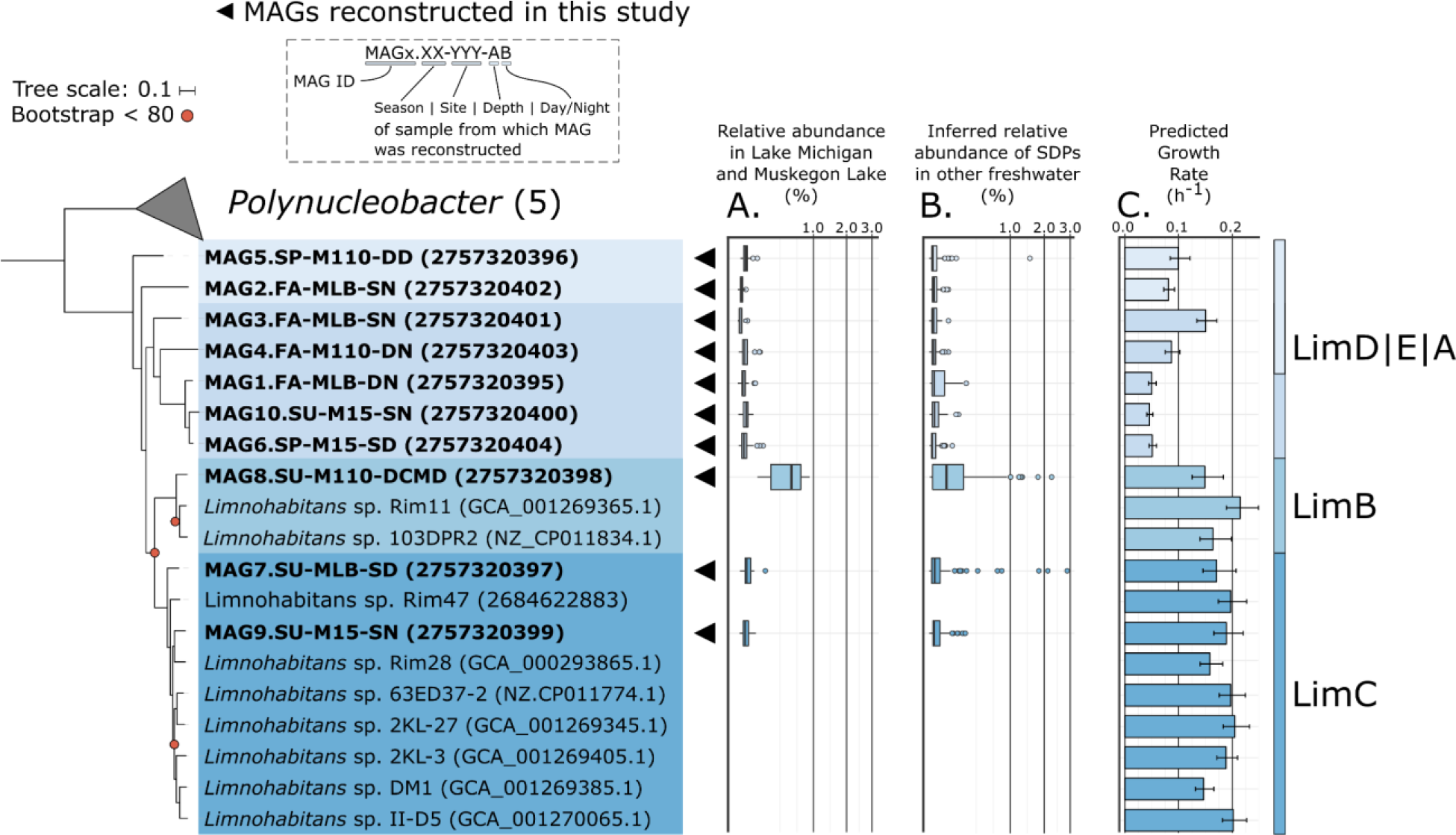
Phylogenomic tree of putative Limnohabitans sp. MAGs and available reference Limnohabitans genomes. MAGs were labeled according to their sample site, season, depth and time of day. Only bootstrap values < 80 are shown. **A.** Boxplots show the normalized relative abundances of each MAG across the 24 samples (square root scale). **B.** Inferred (normalized) relative abundances of closely-related populations to the reconstructed MAGs in 117 freshwater metagenomic datasets publicly available from NCBI (94.5 % identity cut-off). **C.** For each genome or MAG the growth rate was predicted from growth-imprinted genomic traits. The tree was rooted in Chitinophaga niabensis (IMG tax ID: 2636416022).

To assess how prevalent and abundant populations closely-related to our reconstructed MAGs in other freshwater systems are, we competitively recruited reads from 117 publicly-available metagenomic datasets of American and European river, lake and reservoir systems to our set of MAGs (**Figure 2B**). In concordance with previous findings from marker gene surveys, and our Lake Michigan metagenomic survey, populations belonging to the LimB lineage (i.e. closely related to MAG8) are in most freshwater systems the most abundant (43). However, in some systems, populations closely related to other MAGs were also abundant (e.g., MAG7.SU-MLB-SD and MAG1.FA-MLB-DN in Dexter reservoir (OR, USA) and Lake Ontario (ON, CA)); see **Fig. S2 posted at** https://doi.org/10.6084/m9.figshare.10265165 for a detailed description of the inferred *Limnohabitans* population abundances across freshwater environments). Despite its abundance and ubiquity, the underlying functional and genomic adaptations responsible for the success of the LimB lineage remain unclear. The maximum growth rates of the *Limnohabitans* taxa, as predicted from growth-imprinted genome features, suggest the conservation of specific growth rates within currently-defined *Limnohabitans* lineages (**Figure 2C**) (46). Both LimC and LimB lineages had high predicted maximum growth rates, which has been suggested to be a necessary trait to survive protozoan grazing, and to quickly respond to carbon and nutrient pulses (e.g., as a result of phytoplankton blooms) (38, 47).

#### Accessory genomes of Limnohabitans MAGs are enriched in environmental sensing genes

We performed a pangenome analysis using a set of *Limnohabitans* isolate genomes and the Lake Michigan/Muskegon Lake MAGs (n = 19) to pinpoint the functional differences between the abundant LimB MAG8, the other MAGs, and available *Limnohabitans* reference genomes. We observed that most MAGs, even those with high completeness, were missing fragments of the core genome present in the available *Limnohabitans* isolate genomes (**see Fig. S3 posted at** https://doi.org/10.6084/m9.figshare.10265165). This may be the result of genome streamlining (48), evolution under a different ecological setting (49), but likely also the poor recovery of core genome fragments from the metagenomic assembly. We tested for gene set enrichments in the accessory genome of each MAG and found that functional genes involved in environmental sensing (e.g. (ABC-type) transport systems) and secretion systems (e.g., type II and VI secretion systems)) were the categories almost exclusively enriched in five out of ten MAGs and thus may be a differentiator between each MAG’s functional repertoire (**see Fig. S4 posted at** https://doi.org/10.6084/m9.figshare.10265165). The LimC accessory genomes had no significant enrichment in functional gene categories relative to their genome-wide functional categories. The accessory genome of the most abundant MAG8 was composed of 288 genes (~13 % of MAG gene complement). The major difference in the gene content of MAG8 relative to other co-occurring but less abundant MAGs was its specific enrichment in genes for bacterial motility (*pilEF* genes for twitching motility) and chemotaxis (*mcp*, *CheABRW* genes), which were absent from all other MAGs and references genomes except from MAG7 (LimC) and *Limnohabitans* sp. 63ED37-2 (LimB). The *CheABRW* genes were all located on a single 8.6 kb contig (Ga0224460_1280), while flagellar assembly genes (*Flg*, *Flh*, *Fli* gene sets) were present in the *Limnohabitans* core genome enabling active cellular motility. Currently, all available *Limnohabitans* cultures have been phenotyped as non-motile and non-chemotactic. We found no evidence of chemotaxis genes in LimC isolate genomes but did find flagellar assembly pathways in the overall core *Limnohabitans* genome showing that most *Limnohabitans* populations may exhibit motility under certain environmental conditions. Our results thus suggest that the LimB lineage, and possibly also members of the LimC lineage may consist of motile and chemotactic populations that are yet to be cultured. MAG8’s accessory genome was also significantly enriched in high-affinity ABC-type transporters for a variety of (in-)organic components such as iron (III), di- and oligopeptides, polyamines, branched-chain amino acids, phosphate, phosphonate, and sulfate. This suggests that the ability to quickly detect and scavenge resource pulses may confer a competitive advantage to the LimB lineage. A motile lifestyle has also been postulated to play an important role in efficiently scavenging substrates with microscale patchiness, which can be found during events of high primary productivity and *Limnohabitans* abundance (50).

#### Gene expression of Limnohabitans MAG8 across environmental gradients

In conjunction, to gain further insights into what sets the populations represented by MAG8 apart from other less widely distributed and abundant ones, we tested whether the *Limnohabitans* populations represented by the reconstructed MAGs were regulating their overall gene expression differently across the sampled environmental gradients (**Figure 3A**). We assessed changes in gene expression across the spring and fall seasons between which the largest environmental changes occurred (while controlling for sampling location), and between which we thus expected the largest transcriptional responses. We found that compared to all other co-occurring MAGs, MAG8 exhibited a significantly smaller change in gene expression, and this for only a limited number of genes (Benjamini–Hochberg adj. p-value ≤ 0.001, pairwise Wilcoxon rank sum test). Similarly, the differentially expressed genes between the eutrophic (Muskegon Lake) and ultra-oligotrophic station (M110), when controlled for seasonal effects, also showed a smaller change in gene expression for MAG8 than the other MAGs (Benjamini– Hochberg adjusted p-value < 0.001, pairwise Wilcoxon rank sum test). Interestingly, we found that the previously identified chemotaxis genes (*mcp*, *CheAB* genes) were significantly upregulated ([1.1; 3.4] log_2_ fold changes) at the low-nutrient environment of the M110 and M15 sampling sites, suggesting that more active motile behavior is facilitating its ability to thrive at the oligotrophic conditions encountered in Lake Michigan. Phosphorus-dependent chemotaxis has previously been reported to be an adaptation to oligotrophic conditions (51, 52).

**Figure 3:**
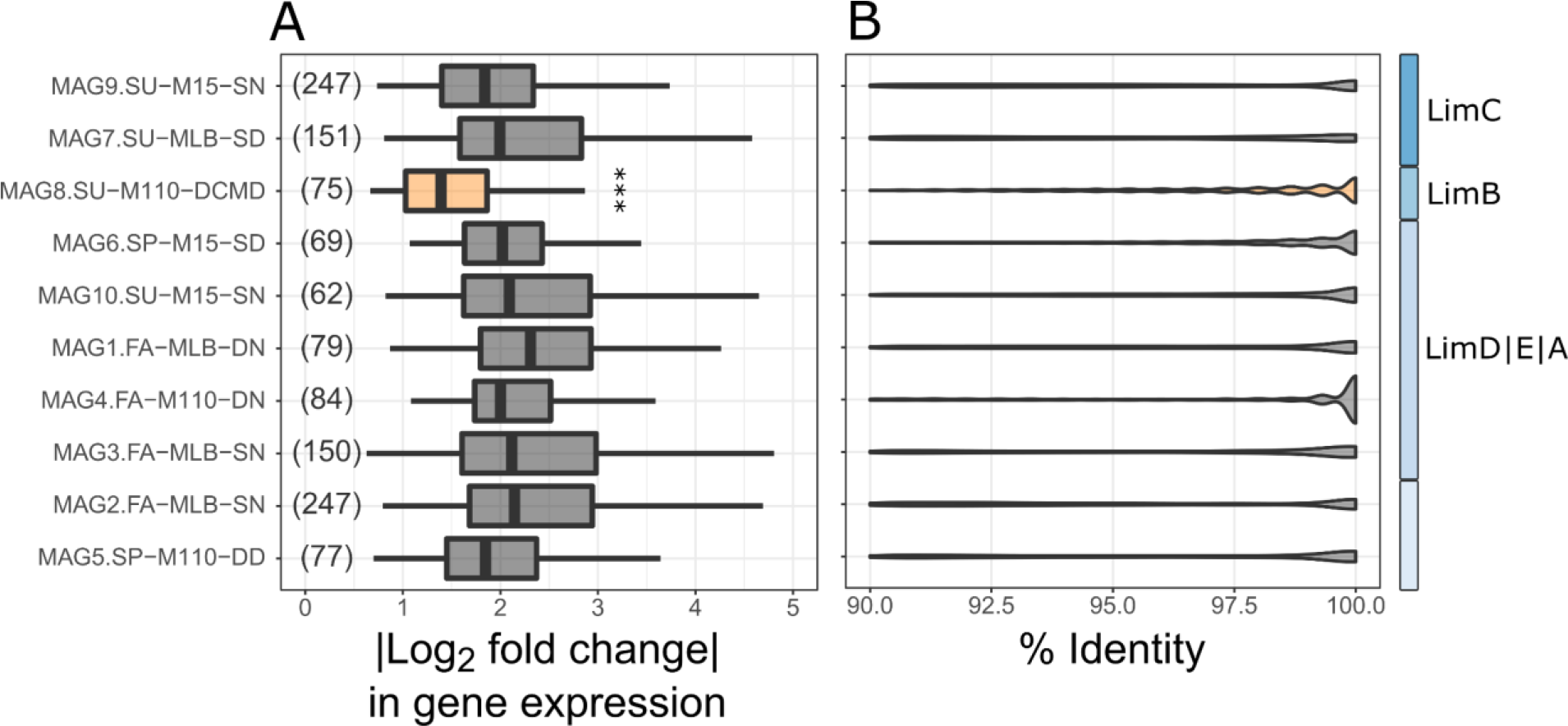
**A.** Absolute values of log_2_ fold changes of differentially expressed genes (adj. p-value < 0.01) between the spring and fall seasons for each MAG (controlled for sampling station). The number of differentially expressed genes of each MAG is indicated between brackets. ***: Pairwise Wilcoxon rank sum test (adj. p-value ≤ 0.001) of MAG8 compared to all other MAGs. **B.** Competitive metagenomic read recruitment to each MAG visualized using violin plots with the bw.nrd0 rule-of-thumb bandwidth. For each MAG the identity profiles of all sample reads were pooled into a single violin plot. The area of each violin plot was fixed, in order to allow visual comparison between MAGs. MAGs were ordered according to their position in the phylogenomic tree.

In addition, we detected 170 differentially expressed genes under differential expression in MAG8 (adj. p-value < 0.01) between the surface and bottom of Lake Michigan station M110 under thermal stratification (summer and fall). The majority of differentially expressed genes of MAG8 were upregulated (n = 134) at the bottom of Lake Michigan. The aerobic carbon monoxide (CO) dehydrogenase medium subunit gene (*CoxM*), was upregulated at the bottom of Lake Michigan, suggesting that energy generation from aerobic CO oxidation (i.e. methylovory) may be an important trait in this environment. The source of CO in the deep regions of Lake Michigan is unknown but could be a result of the incomplete decomposition of humic acids and phenolic compounds in the sediment (53). Our findings regarding C1 oxidation expression are in line with the expression patterns of a Chloroflexi population along the water column at the offshore site in Lake Michigan (54). The expression of several high-affinity branched-chain amino acid genes was upregulated. Although this may indicate that a shift in amino acid preference to branched-chain amino acids could be important, laboratory experiments, which although limited in taxonomic resolution, found that amino-acid specificity is rather rare in freshwater taxa (55). In combination with the lower (in-)organic transporter gene expression for phosphonate, iron(III), and sulfate these results strongly suggest a lower (in-)organic nutrient uptake by the LimB lineage at the bottom of Lake Michigan.

### Genomic microdiversity

#### Sequence Discrete Populations of *Limnohabitans*

MAGs represent consensus population genomes and can encompass additional strain-level variation (10, 13, 56). In addition to testing the level of functional differentiation between the MAGs, we screened the MAGs for the potential presence of microdiversity based on their metagenomic recruitment profiles (**Figure 3B;** detailed recruitment profiles are provided in **Fig. S5 and S6 posted at** https://doi.org/10.6084/m9.figshare.10265165). The presence of distinct local maxima at < ~99 % identity indicated the presence of sequence variants in the metagenomic data for which we were unable to reconstruct the genome and are commonly referred to as Sequence Discrete Populations (SDPs) (56, 57). Several MAGs appeared to have multiple SDPs but the strongest support was found in MAG8, which was congruent with the large number of reconstructed 16S rRNA genes for the LimB lineage. In almost every sample we found SDPs of MAG8 confined to the 90 % – 97 % nucleotide identity range (**see Fig. S5 posted at** https://doi.org/10.6084/m9.figshare.10265165). The Muskegon Lake site, which was the most diverging environment in terms of nutrient levels, had primarily read recruitment at < 99 % identity suggesting that in this environment more evolutionary distant LimB populations were present from the one represented by MAG8. In addition, this microdiversity may explain the difficulty in fully assembling this high-coverage MAG as many assembly tools struggle in resolving the genomes from related strains from metagenomic data (58). In our case, normalizing to a lower and more even sample coverage, or downsampling to a fixed number of reads did not improve the assembly of MAG8, again suggesting that microdiversity and not excessive coverage as the cause of the poor assembly. The presence of SDPs at various sites and during different seasons indicated the presence of microdiversity in this MAG. However, care must be taken with the interpretation of these density profiles, as the level of smoothing dictated by the bandwidth parameter can either overemphasize spurious local maxima (undersmoothing), or instead confound closely related populations (oversmoothing) (59). Therefore we used, and recommend other researchers to only use these profiles as qualitative indicators for the presence of microdiversity and/or closely-related populations.

#### Microdiversification in *Limnohabitans* MAGs across environmental gradients

In order to determine whether these SDPs were actually indicative of the presence of different strains we used the DESMAN workflow to infer the number of strains and their abundance based on variant base frequencies and coverage patterns of 36 single copy core genes (12). A total of 23 strains could be resolved from the eight initial putative Limnohabitans MAGs (**Table 2**), with most MAGs representing at least two strains, and MAG8 representing a total of six individual strains. Within each MAG, the strains exhibited both positive and negative correlations with environmental parameters previously associated with *Limnohabitans* marker-gene based subtype abundances (**Figure 4A**) (43). Several strain frequencies were strongly correlated to the total phosphorus concentration, but the majority showed strong correlations with water temperature, and thus indirectly with seasonality and sampling site as well. Light availability, as measured by photosynthetically active radiation (PAR), did not strongly correlate with strain frequencies. Two MAGs of the LimDEA lineages (MAG4, MAG10), two MAGs of the LimC lineage (MAG7), and the LimB MAG (MAG8) had strains that were correlated either positively or negatively with the water temperature. Only strains within the LimB and LimC lineages showed strong correlations with the total phosphorus concentration in the water. In case of the LimA lineage, preferences towards higher water temperature and specific carbon sources (i.e. allochthonous DOM) have previously been shown (32), which is congruent with our observations on the LimDEA inferred strain abundance.

**Figure 4:**
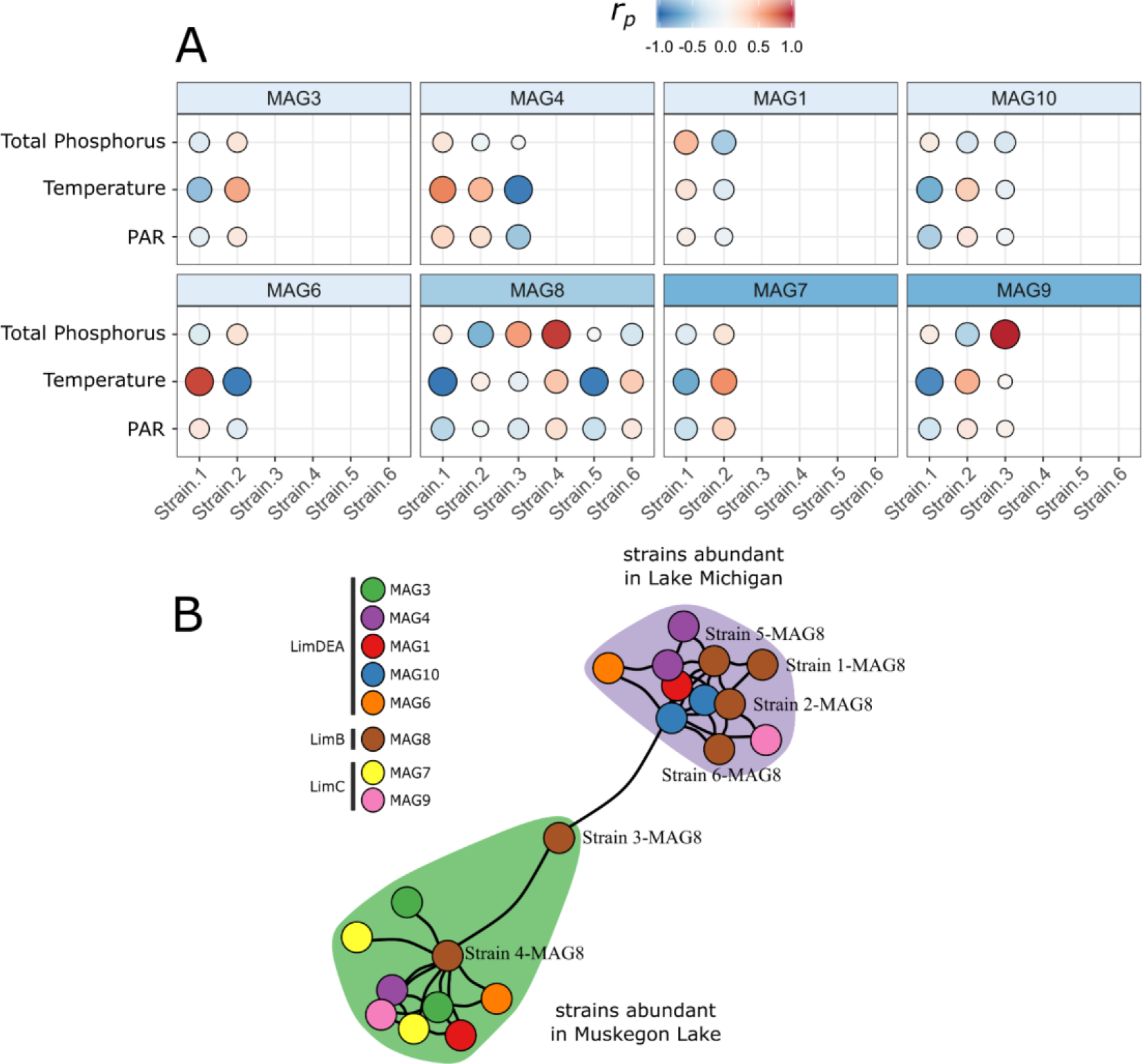
**A.** Pearson’s correlation (r_P_) between Limnohabitans MAG frequencies and primary environmental parameters differentiating the environmental gradients of the studied transect. Size of labels is proportional to correlation strength. PAR: photosynthetically active radiation **B.** Co-occurrence network of all Limnohabitans strains using a Fruchterman-Reingold layout. Colored regions highlight identified network modules (n = 24).

Using network analysis, we found that the delineated strains clustered into two distinct modules of an approximately equal number of co-occurring strains (**Figure 4B**). One network module contained primarily strains with seasonal frequency increase in Muskegon Lake, while the other contained strains with seasonal frequency increase in Lake Michigan. The differences in environmental conditions associated with these sampling sites (e.g. trophic state and temperature) appear to have given rise to the existence of two sub-communities of *Limnohabitans* populations with correlating seasonal abundance profiles. Certain MAGs only had strains in a single network module showing that environmental specificity may already be dictated at the MAG-level, such as Muskegon Lake for MAG7 and Lake Michigan for MAGs 3 and 10. The other MAGs had a more balanced distribution over the network modules, and MAG8 had the majority of its strains in the Lake Michigan associated network module.

Focusing on the strain diversity of MAG8, we found that it was strongly associated with the spatiotemporal components of the data (**Figure 5**). The six inferred LimB strains displayed strong environmental specificity, with two strains dominating under eutrophic conditions found in the Muskegon Lake watershed, and four other strains dominating across the more oligotrophic Lake Michigan transect. Based on the SDP analysis the strains abundant in Muskegon Lake appeared to be the most evolutionary distant from the reconstructed consensus MAG (i.e. majority of read recruitment at ≤ 95 % identity; **see Fig. S5 posted at** https://doi.org/10.6084/m9.figshare.10265165). This highlights that the largest evolutionary distance existed between the set of strains adapted to different trophic conditions found in Lake Michigan and Muskegon Lake. In addition, all strains had clearly distinct seasonal abundance profiles that were either positively or negatively correlated with the season. For example, five strains were inferred to be present in the spring at both the surface and the deepest region of the oligotrophic M110 site where temperature and overall environmental conditions remain stable. During the summer and fall seasons, five out of the six strains were still present at deepest region of the oligotrophic M110 site, while only three remained at the surface (see **Fig. S7 posted at** https://doi.org/10.6084/m9.figshare.10265165).

**Figure 5:**
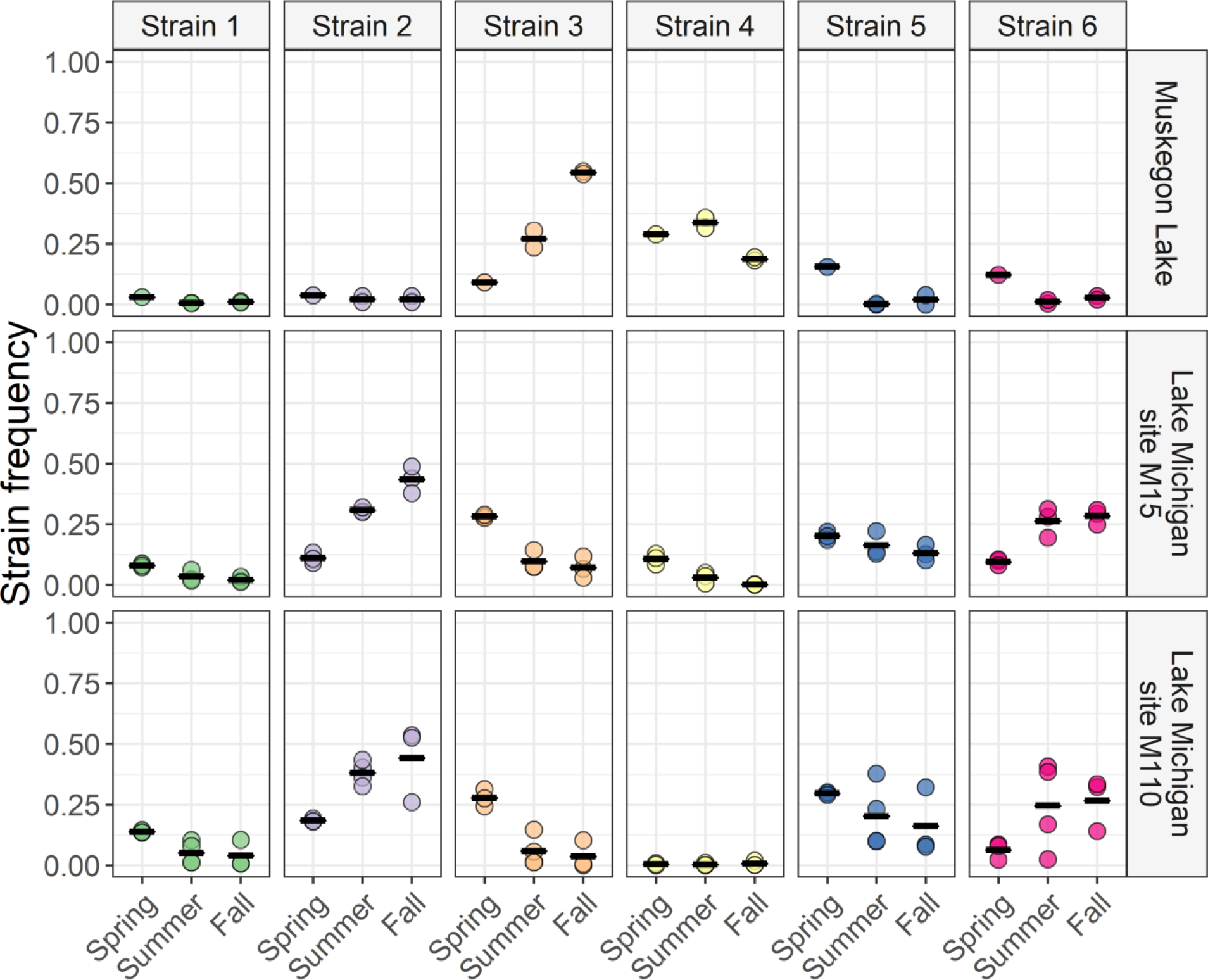
Spatiotemporal strain frequency dynamics of MAG8.SU-M110-DCMD as inferred from DESMAN. Only strains for which inference was robust are shown, and mean frequencies per site and season are indicated by horizontal bars.

Overall, 16.1 % of the variation in strain composition could be explained by the season (spring, summer or fall), with an additional 25.4% of seasonal variation being conditional on the sampling site (**see Fig. S8 posted at** https://doi.org/10.6084/m9.figshare.10265165). 12.6 % of the variation was explained solely by the sampling site. The strain alpha diversity at the offshore oligotrophic M110 sampling station was among the highest measured, and unlike other sampling locations, largely invariant to seasonality (**see Fig. S9 posted at** https://doi.org/10.6084/m9.figshare.10265165). While this is in concordance with the relatively stable environmental conditions at the M110 site, such a broad abundance distribution across depth and nutrient gradients has not yet been reported for other abundant freshwater taxa (e.g., Actinobacteria (60), *Polynucleobacter (5, 61)*). Data from extensive marker gene surveys have shown that other *Limnohabitans* taxa, such as those in the LimA lineage have a preference for surface waters but can be found at greater depths as well (32, 42). The alpha diversity of the strains was highest in spring and lowest in the fall (approximately 50 % reduction). The maximum diversity during spring is in line with previous studies which have reported increased levels of microdiversification during peaks of primary production (43), and higher community diversity levels that have been attributed to the increased resource heterogeneity that may occur early in the season (28). Strain diversity of LimB, and also the other MAGs, appeared to follow the overall taxonomic diversity patterns across the seasons.

In conclusion, we investigated the genomic and functional diversity of the ubiquitous and prevalent *Limnohabitans* taxa across a transect of Lake Michigan, part of the largest freshwater ecosystem in the world. Our findings show that thermal adaptation may be a more important driver of overall microdiversification within the *Limnohabitans* genus, but that specifically for the LimB and LimC lineages, microdiversification under influence of nutrient availability is an additional driver that is associated with significantly larger genomic divergence. These findings are corroborated by previous work on *Vibrio* and other *Limnohabitans* taxa showing that closely related strains can have remarkably different environmental adaptations (6, 38, 43). However, due to the strong correlation of environmental conditions and sampling sites described in this study, additional supporting studies with more extensive environmental data will be necessary to avoid missing other potential drivers of microdiversity. Lineage-level expression analysis indicated that, compared to other *Limnohabitans* lineages, LimB displayed smaller shifts in gene expression across the sampled gradients, possibly due to the genomic heterogeneity in strains optimized to specific conditions, and that a shift to methylovory and increased chemotaxis were possible adaptations for handling steep depth and/or nutrient gradients. The *Limnohabitans* genus has been proposed to be equivalent in importance for freshwater food webs, as is the SAR11 taxon for marine food webs (43). While marker-gene surveys have learned us a great deal about this important group of bacterioplankton, SAR11 has arguably been studied more intensely on its genomic, functional and evolutionary aspects. Recent genome-base efforts on *Limnohabitans* have just started to uncover the vast metabolic portfolio of this genus (35). Future *in situ* metagenomics, targeted cultivation efforts and synthetic ecology experiments on this genus are thus needed to enable a detailed understanding of the functional, phenotypic and ecological implications that are associated with its microdiversification.

## Materials and methods

### Metagenomic and metatranscriptomic data

We used Lake Michigan and Muskegon Lake publicly available metagenomic and metatranscriptomic data of the 0.22- to 3-μm plankton fraction that were previously collected at three stations (i.e. labeled MLB, M15, M110) spanning a productivity gradient (28). These samples were taken at three time points, divided over three seasons (i.e. spring, summer, fall of 2013), at different depths (5m up to 108m), and time of day (day/night). Sampling and DNA/RNA extraction of these samples have been described elsewhere (54). Libraries were prepared using the Nextera (summer and fall samples) or TruSeq (spring samples and all metatranscriptomic samples) preparation kits (Illumina Inc.), and sequenced on a 2 × 150 bp paired-end HiSeq 2000 sequencing system. *Sickle* (v1.33.6; https://github.com/najoshi/sickle) was used for removing erroneous and low-quality reads from the data. *Scythe* (v0.993; https://github.com/vsbuffalo/scythe) was used for removing adapter contaminant sequences. Denoised reads were evaluated using FastQC.

### 16S rRNA gene reconstruction

We reconstructed full-length 16S rRNA gene sequences from quality-trimmed reads using *EMIRGE* (v0.60.3) (62). *EMIRGE* was run using the non-redundant 97%-clustered Silva database (v123; (63)) as reference. *EMIRGE* was run for 65 iterations with the quality-trimmed reads and with an insert size of 200 and a standard deviation of 50, as these were known parameters. The *EMIRGE* reconstructed sequences were dereplicated and sequences that fell outside the 1,000 bp - 1,700 bp range or contained ambiguous bases were discarded. We classified the sequences according to the *Limnohabitans* specific framework by Kasalický *et al.* (2013) by evaluating their phylogenetic placement relative to the reference sequences, and by the freshwater classification framework by Newton *et al.* (2011) through the *TaxAss* pipeline (27, 38, 64). Classification of the sequences using the *TaxAss* pipeline combines both the Silva v123 database and a manually-curated freshwater taxonomy database (FWDB) (64). FWDB classification was favored over the Silva classification if the length-corrected identity to a FWDB sequence was > 97 % (64). Sequences were clustered into OTUs at an average similarity of 97% using the average nearest neighbor method in *mothur* (v1.37) (65).

### Genome reconstruction

The reads were dereplicated using a custom perl script and interleaved into a single sequence file for subsequent assembly (66). Interleaved sequences were digitally normalized using *bbnorm* to a target 60× coverage and reads with a coverage less than 5× were discarded in order to improve the subsequent assembly step (67, 68). The interleaved reads of all 24 samples were assembled on a per-sample basis into contigs using *IDBA-UD* (v1.1.3) with kmer lengths varying from 41 to 101 in steps of 10 (69). Contigs were taxonomically classified at the order level by a diamond search against the non-redundant NCBI protein database using *DESMAN* scripts (12) (*classify_contigNR.pl* with MIN_FRACTION = 0.1), after which the classification output files were formatted into an annotation file compatible with the *Vizbin* binning tool. Initial Metagenome Assembled Genomes (MAGs) of all classified Betaproteobacteria were retrieved through a manual binning strategy in *Vizbin* (v0.9, default settings, minimum contig length = 2,000 bp) (70). Quality-trimmed reads were mapped to each individual assembly using *bwa-mem* (v0.7.8) on default settings (71). *Samtools* (v1.3.1) was used to convert, sort and index the sam-files (72). From this initial set of 92 population genomes we extracted the unique representative genomes by taking into account the Average Nucleotide Identity (*pyani* v0.2.7 – ANIb method) between MAGs as well as the completeness statistics inferred from a CheckM analysis (v1.07). If the ANI was > 99 % between genomes and the redundancy <10%, the MAG with the highest completeness was chosen as representative. This approach reduced the number of MAGs from 92 betaproteobacterium MAGs to 10 putative *Limnohabitans* MAGs. Next, all sample reads were competitively mapped to the putative *Limnohabitans* MAGs. The *Anvi’o* platform (v2.3.0) was then used to manually refine the unique MAGs identified through *Vizbin* by evaluating differential coverage patterns across the samples. Completeness and redundancy estimates of the final MAGs were estimated by *CheckM* (v1.07) (73). Mean relative MAG abundances, normalized for observed genome size, were calculated from the mean coverages exported from *Anvi’o* as follows:

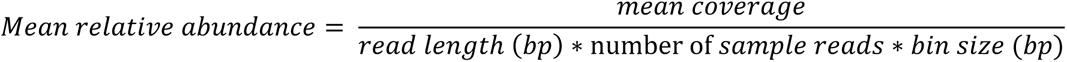

The presence of closely related populations (sequence discrete populations, SDPs) in publicly available datasets (n = 117) used in Neuenschwander et. al (2017) was assessed by blasting one million reads of each dataset to the 10 putative *Limnohabitans* MAGs and assessing the read alignment across the contigs of the MAGs (60). The relative abundance of SDPs was inferred by normalizing the number of reads that aligned with > 94.5% identity, with the genome sizes of the corresponding MAGs (in Mbp), and the total number of reads mapped (i.e. one million).

The refined MAGs were submitted for gene-calling and annotation to the Joint Genome Institute’s Integrated Microbial Genomes isolate annotation pipeline (74). For each MAG and selected publicly available freshwater genomes the minimal generation time (MGT) and optimal growth temperature were predicted based on “growth-imprinted” genome features by means of the Growthpred (v1.07) software (46). The predicted specific growth rates (*μ*_*spec*_) were then calculated by the following formula:

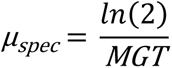

Additionally we calculated the index of replication (*iRep*), a measure for the population-averaged replication status at the time of sampling, for each MAG (45). Sequence discrete populations were examined by competitively mapping one million reads per sample (*blastn* v2.25; (75)) and evaluating the nucleotide identity profiles of the best hits for each MAG.

### Phylogenetic and phylogenomic tree construction

A 16S rRNA gene phylogenetic tree was constructed using a set of publicly available betI-clade, *Limnohabitans* genus, and other related lineage sequences. Sequences were aligned using the SINA aligner (76). The tree was constructed with *Fasttree* (v2.1.9) using the GTR+CAT evolutionary model and run in sensitive mode (−spr 4 −mlacc 2 –slownni options) (77). The phylogenomic tree was constructed based on the codon alignment of a set of 37 conserved marker genes through *Phylosift* (78). The tree was generated from the codon alignment by means of *RAxML* (v8.2.8) with the GTRGAMMA model and 1,000 bootstraps (79). Both trees were visualized and annotated in *iTOL* (80), exported and further annotated in *Inkscape* (v0.91).

### Strain inference

We used the standard workflow of the DESMAN software for (i) inferring strain frequencies and (ii) assigning accessory genomes to robust strains in the MAG of interest (12). Briefly, gene calling was performed with *prodigal* (v2.6.3) (81), and single copy core genes (SCCGs) were detected by means of reversed position specific blast (*RPS-blast*) against a SCCG *E. coli* reference database provided with DESMAN. Variant positions were then found using the base count frequencies of these SCCGs with the *Variant_filter.py* script (*-p* for 1d optimization of minor variant frequency detection). Samples were required to have a coverage > 1× (*−m*). SCCGs were filtered based on their median coverage (*−c*) and a maximum divergence of 2.5 on the median coverage was allowed for each SCCG. The initial coverage cut-off was set at 25.0 (*−f* flag) and SCCGs were filtered if less than 80% of the samples passed the coverage filter (*−sf* flag). The identified variant positions were then used as input for the DESMAN haplotype inference tool. We ran 10 replicate runs of DESMAN for up to 12 haplotypes. The final robust number of strains was determined using the *resolvenhap.py* script, and for which the average single nucleotide variation errors were < 10 % (**see Table S2 Posted at** https://doi.org/10.6084/m9.figshare.10265165). For inference on the abundance of each inferred strain, the Gamma output files were used. The scripts used to perform this analysis are provided at: https://github.com/rprops/MetaG_analysis_workflow/wiki/21.-DESMAN.

### Network analysis

Co-occurrence networks on the Limnohabitans inferred strains were inferred using *SparCC* as implemented in the *SpiecEasi* package (v1.0.6) (82, 83). The input data consisted of the strain frequency data multiplied with the normalized abundance of each MAG. 100 iterations were run on both the outer and inner loop, and a correlation threshold of 0.2 on the Pearson’s correlation values was put on the inner loop. The graphs were visualized using the *igraph* package (v1.2.4.1) for all correlations with an absolute value greater than 0.3 and empty nodes were discarded. Network modules were identified via a spin-glass model and simulated annealing as implemented in *igraph* under default settings.

### Expression analysis

Denoised metatranscriptomic reads were competitively mapped to the set of unique Betaproteobacteria MAGs. Testing for differential expression of genes in the *Limnohabitans* MAGs was performed with *DESeq2* (v1.18.1) on default settings (84). Genes were considered under differential expression if their Benjamin-Hochberg corrected P-value was < 0.01.

### Pangenome analysis

We applied the default pangenome analysis workflow available in *Anvi’o* (v2.3.0) to all *Limnohabitans* genomes (85). Gene calling was performed with *prodigal* (v2.6.2), amino acid sequences were aligned with *muscle* (v3.8.31) (86), amino acid similarities calculated with *blastp* (v2.2.29), and sequences were clustered with *mcl* (v14-137) (87). Accessory genomes were manually binned in *Anvi’o* (**see Fig. S3 posted at** https://doi.org/10.6084/m9.figshare.10265165). As there were no up to date IMG annotation projects available for the other *Limnohabitans* genomes, the gene calls from *Anvi’o* were exported and annotated with KEGG orthology identifiers using the BlastKOALA web server (taxid: 665874, species_prokaryotes database) to facilitate direct comparison between genomes (88). Enrichment of functional categories (e.g. KEGG subsystems) was conducted with the hypergeometric testing available in the *clusterProfiler* package (v3.6.0) (89). P-values were adjusted for multiple testing with the Benjamini-Hochberg correction.

## Availability of data and material

Supplementary information is available at https://doi.org/10.6084/m9.figshare.10265165. Raw sequence reads are available from the JGI portal under the project IDs specified in **Table S1** posted at https://doi.org/10.6084/m9.figshare.10265165. MAGs and their genome annotations are available from IMG under the taxon OIDs specified in **Table 1**. The data analysis workflow is publicly available at https://github.com/rprops/Limno_lakeMI. Pangenome analysis input and output files are available from https://doi.org/10.6084/m9.figshare.7547159. The Anvi’o profile database (v2.3.0) used to refine the MAGs is available from https://doi.org/10.6084/m9.figshare.7547852.

## Acknowledgements

We thank the crew on the R/V Laurentian and Ann McCarthy and Marian Schmidt for sampling of Lake Michigan and Muskegon Lake, and Ann McCarthy for DNA and RNA extractions. Part of this work was supported by funding to V.J.D. by the Community Sequencing Program (U.S. Department of Energy Joint Genome Institute, a DOE Office of Science User Facility, supported under Contract No. DE-AC02-05CH11231), the National Science Foundation (Award No. 1737680), and the University of Michigan. R.P. was supported by Ghent University (BOFDOC2015000601) and a Sofina Gustave-Boël grant from the Belgian American Educational Foundation (BAEF).

